# Altered striosome-matrix distribution and activity of striatal cholinergic interneurons in a model of autism-linked repetitive behaviors

**DOI:** 10.1101/2024.05.23.595498

**Authors:** Jordan Molitor, Juliette Graniou, Pascal Salin, Francis Castets, Ahmed Fatmi, Lydia Kerkerian-Le Goff, Laurent Fasano, Xavier Caubit, Paolo Gubellini

**Author notes:** Contributed equally (Jordan Molitor, Juliette Graniou, Pascal Salin).

## Abstract

Repetitive behaviors are cardinal features of many brain disorders, including autism spectrum disorder (ASD). We previously associated dysfunction of striatal cholinergic interneurons (SCINs) with repetitive behaviors in a mouse model based on conditional deletion of the ASD-related gene *Tshz3* in cholinergic neurons (*Chat-cKO*). Here, we provide evidence linking SCIN abnormalities to the unique organization of the striatum into striosome and matrix compartments, whose imbalances are implicated in several pathological conditions. *Chat-cKO* mice exhibit altered relationship between the embryonic birthdate of SCINs and their adult striosome-matrix distribution, leading to an increased proportion of striosomal SCINs. In addition, the ratio of striosomal SCINs with slow-irregular *vs*. sustained-regular firing is increased, which translates into decreased activity, further stressing the striosome-matrix imbalance. These findings provide novel insights onto the pathogenesis of ASD-related stereotyped behaviors by pointing to abnormal developmental compartmentalization and activity of SCINs as a substrate.

## INTRODUCTION

Autism spectrum disorder (ASD) is characterized by abnormalities in two core behavioral domains, namely deficits in social interactions and restrictive, repetitive patterns of behavior, also known as stereotypies (DSM-5, 2013). Clinical and animal studies have linked dysfunction of the striatum, the main input station of the basal ganglia, to the pathogenesis of ASD and pointed to the corticostriatal pathway and to striatal spiny projection neurons (SSPNs) as main players (Fuccillo, 2016; Li and Pozzo-Miller, 2019). Moreover, in recent years, the role of cortical and striatal interneurons in ASD is attracting increasing interest (Contractor et al., 2021; Rapanelli et al., 2017a). Pathological repetitive behaviors occur as well in other brain disorders, including Tourette’s syndrome, obsessive-compulsive disorder and schizophrenia, and can be driven by drugs of abuse (Figee et al., 2016; Gerdeman et al., 2003; Jutla et al., 2022; Ridley, 1994). In this context, there is growing evidence from studies in human and rodent models linking repetitive behavior to the loss or dysfunction of striatal cholinergic interneurons (SCINs) (Aliane et al., 2011; Crittenden et al., 2017; Kataoka et al., 2010; Lennington et al., 2016; Martos et al., 2017; Rapanelli et al., 2017a; Xu et al., 2021; Xu et al., 2015). Despite their small number, SCINs have morphofunctional features that place them as key modulators of striatal microcircuits, playing a crucial role in habit-mediated and goal-directed behavior as well as in attentional set-shifting, conferring behavioral flexibility (Prado et al., 2017). SCINs are tonically-active neurons with a giant cell body and an extended neuritic arborization at the interface between the striatal input systems and SSPNs (Abudukeyoumu et al., 2019; Ahmed et al., 2019; Apicella, 2017; Calabresi et al., 2000; Goldberg and Wilson, 2010). In addition, they are considered as leading players in the communication between the two intermingled compartments that confer a unique mosaic structure to the striatum, namely the striosomes (also called patches) and the matrix, due to their neurites crossing the borders of the two compartments. The striosomes and matrix differ in their neurochemical features, birthdate of their populating neurons, input-output connections (in particular their cortical inputs from limbic and sensorimotor areas), and have presumably distinctive functional attributes in a range of behaviors and pathological conditions (Brimblecombe and Cragg, 2017; Crittenden and Graybiel, 2011; Gerfen, 1992; Graybiel and Matsushima, 2023; Kawaguchi, 1997). Recent work reported altered striosome-matrix compartmentalization in ASD patients (Kuo and Liu, 2020) and in an ASD mouse model (Kuo and Liu, 2017), and suggested a preferential involvement of the striosomal compartment in the context of drug-induced stereotypy (Murray et al., 2015). Despite the recognized role of SCINs in pathological repetitive behaviors, the relationship between dysfunction of these interneurons, their striatal compartmentalization and repetitive behaviors has not been investigated.

Here we addressed this issue in the context of ASD. We previously identified *TSHZ3* (teashirt zinc-finger homeobox family member 3, also known as *ZFP537*) as an ASD-related gene, and provided evidence that cortical projection neurons and SCINs are main determinants of the ASD-like behavioral abnormalities linked to *Tshz3* deletion in mouse models (Caubit et al., 2021; Caubit et al., 2016; Caubit et al., 2022). TSHZ3 is a transcription factor regulating the expression of numerous genes involved in brain development and functioning (Caubit et al., 2021; Caubit et al., 2016; Chabbert et al., 2019; Kang et al., 2011). In the cortex, TSHZ3 is expressed in most cortical projection neurons and in ∼30% of GABAergic interneurons. In the striatum, TSHZ3 is not expressed in SSPNs but in virtually all SCINs, which represent more than 90% of the TSHZ3-expressing cells in this structure. In contrast, TSHZ3 is not expressed or is expressed in a low proportion of neurons in the other main cholinergic systems of the brain (Caubit et al., 2022). Conditional deletion (cKO) of *Tshz3* in cortical projection neurons (*Emx1-cKO* mice) or in SCINs (*Chat-cKO* mice) does not modify the number of these neurons but alters, respectively, their synaptic function and their electrophysiological features. Moreover, *Emx1-cKO* mice show impaired social interactions, while *Chat-cKO* show repetitive/stereotyped behaviors, thus segregating the two ASD behavioral domains (Caubit et al., 2022; Roubertoux et al., 2020). Altogether, these data identify SCINs as key actors in the stereotyped behaviors characterizing *TSHZ3*-linked ASD. In this study, we investigated the pathological changes in the developmental trajectory and electrophysiological properties of SCINs in relation to their striosome *vs*. matrix compartmental distribution in the context of *Tshz3*-linked ASD, as an entry point to unveil novel neuronal substrates and mechanisms underlying repetitive/stereotyped behaviors.

## RESULTS

### Deletion of *Tshz3* from committed postmitotic cholinergic neurons of the striatal anlage in *Chat-cKO* mice

As a first step to study SCIN developmental processes, and how and when they are affected by *Tshz3* deletion, we characterized the time-course of the expression of cholinergic markers and of *Tshz3* in the developing mouse brain. The generation of SCIN progenitors occurs in mouse between embryonic day (E) 12 and E15 in the medial ganglionic eminence (MGE), septal neuroepitelium and preoptic area (Knowles et al., 2021; Marin et al., 2000; Pappas et al., 2018), and choline acetyltransferase (*Chat*) mRNA expression has been reported at E14.5 in the post-mitotic zone of the lateral ganglionic eminence (LGE), which corresponds to the striatal anlage (Furusho et al., 2006; Lopes et al., 2012). We previously showed that *Tshz3* mRNAs are detected as early as E12.5 in the LGE and MGE (Caubit et al., 2005). Here, we identified cholinergic cells using ChAT and vesicular acetylcholine transporter (VAChT) immunostaining or by tdTomato fluorescence in *Chat-Cre;Ai14* mice. To identify TSHZ3-expressing cells, we performed immunostaining of either TSHZ3 (in *Chat-Cre;Ai14* mice) or ß-Gal (in *Tshz3^+/lacZ^* mice), which both provide nuclear labeling (Caubit et al., 2021; Caubit et al., 2016). Interestingly, at E13.5, expression of VAChT and ß-Gal is detected at the level of the striatal anlage, with some cells co-expressing both markers (Fig. 1a), while no labelling is obtained with anti-ChAT and anti-TSHZ3 antibodies. At E14.5, most cells of the striatal anlage co-express VAChT and ß-Gal (Fig. 1b), but ChAT is still not detectable by immunostaining. At the same stage, TSHZ3 is detected in the tdTomato-positive cholinergic cells, showing that the *Chat-Cre* is active during embryogenesis in committed, immature cholinergic neurons of the striatal anlage (Fig. 1b). Finally, at E17.5, most TSHZ3-positive cells in the immature striatum co-express ChAT (Fig. 1c), similarly to the adult (Caubit et al., 2021; Caubit et al., 2022), as well as VAChT (Fig. 1c). At this time point, the number of TSHZ3-positive cells is drastically reduced in *Chat-cKO* mice compared to control (Fig. 1d), demonstrating the embryonic loss of TSHZ3 expression. Overall, these data indicate that VAChT is an earlier marker of SCINs than ChAT, and that the *Chat-Cre*-mediated deletion of *Tshz3* occurs in cholinergic cells from E14.5 onwards.

**Fig. 1.**
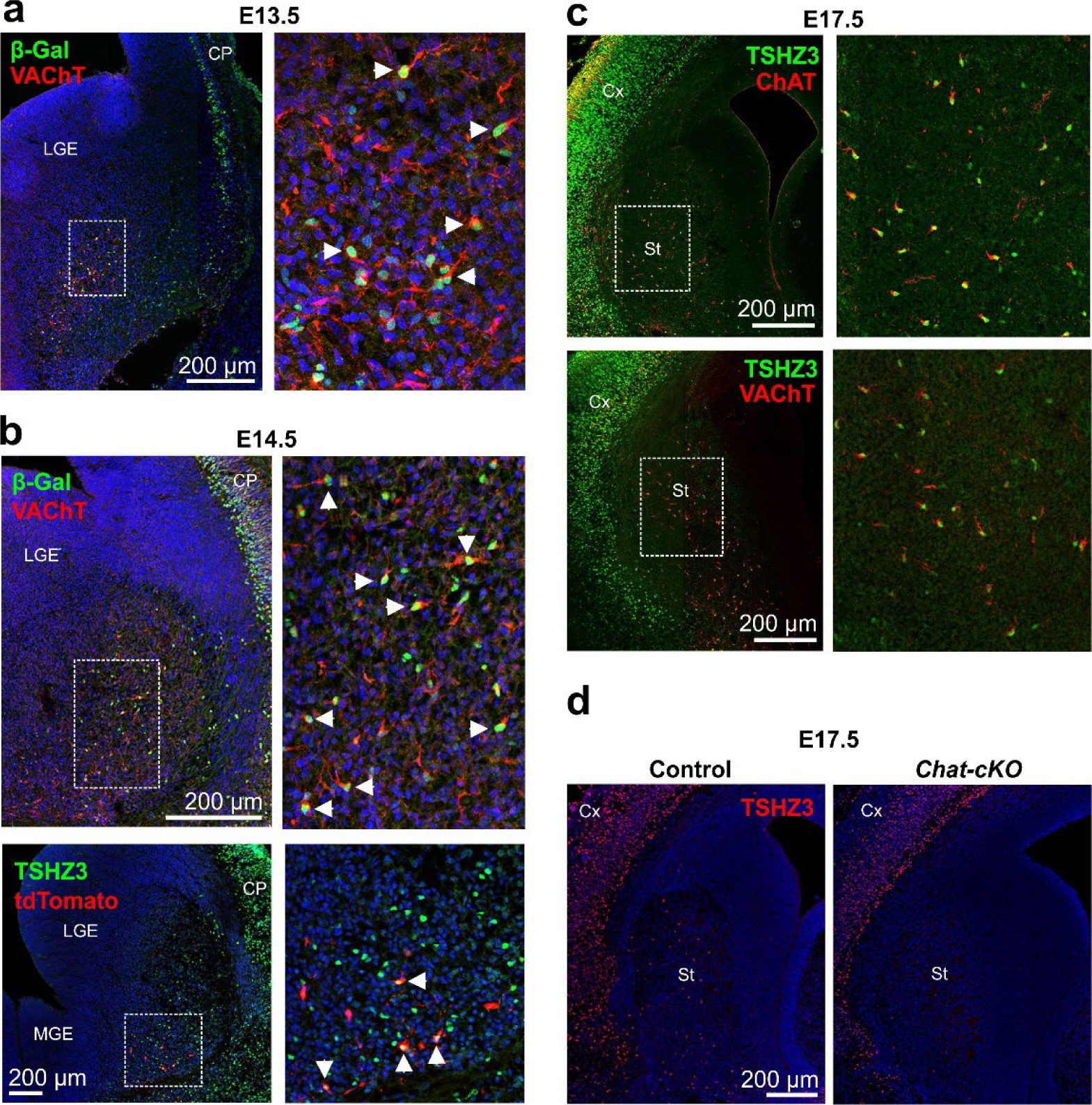
Expression of cholinergic markers and *Tshz3* in the embryonic mouse brain and *Tshz3* deletion in *Chat-cKO* mice. **a**, Coronal section of an E13.5 *Tshz3^+/lacZ^*embryo showing ß-Gal-positive (i.e., expressing *Tshz3*, green) and VAChT-positive (red) cells forming a cluster in the ventrolateral part of the basal forebrain (striatal anlage). The inset is a maximum intensity projection (MIP) of 21 xy images with an axial distance of 0.42 µm between planes, showing that several of these cells co-express ß-Gal and VAChT (arrowheads). **b**, Coronal sections at E14.5 in which *Tshz3* expression is detected in the striatal anlage by ß-Gal (upper row, green) or TSHZ3 immunostaining (lower row, green) and cholinergic cells are revealed by VAChT immunostaining (upper row, red) or tdTomato fluorescence (lower row, red). Upper images are from a *Tshz3^+/lacZ^*mouse, lower images from a control (*Chat-Cre;Ai14*) mouse. Right panels show that most cholinergic cells co-express *Tshz3* (arrowheads). **c**, Dual immunodetection of ChAT or VAChT and TSHZ3 on coronal sections of the embryonic striatum at E17.5. Note that almost all cholinergic cells are TSHZ3-positive (see insets). Images are from wild-type mice. Right panels show MIPs of 4 xy images with an axial distance of 0.69 µm between planes. **d**, Coronal sections at E17.5 showing the drastic reduction of TSHZ3-positive cells (red) in the striatum of a *Chat-cKO* mice compared to control. In **a**, **b** and **d** sections are counterstained with DAPI (blue). Abbreviations: CP = cortical plate; Cx = cortex; LGE = lateral ganglionic eminence; MGE = medial ganglionic eminence; St = striatum.

### Altered compartmental distribution of late-born SCINs in *Chat-cKO* mice

We then tested whether *Tshz3* deletion affects SCIN neurogenesis in relation to their striosome-matrix distribution in the adult striatum. Previous studies in the rat showed that early-born SCINs populate preferentially the striosomes and late-born SCINs the matrix (van Vulpen and van der Kooy, 1998). However, the relative birthdate/compartment relationship for SCINs has not been established in the mouse. To address this question, we performed birth-dating experiments by injecting 5-ethynyl-2′-deoxyuridine (EdU) in pregnant control and *Chat-cKO* mice at E10.5, E11.5, E12.5, E13.5 or E14.5, and µ-opioid receptor 1 (MOR1) immunostaining to identify striosomes. Neuronal precursors differentiating into SCINs after these injections were then visualized in MOR1-positive striosomes or MOR1-negative matrix of adult offspring mice by ChAT immunostaining and EdU staining (Fig. 2a). Thanks to this approach, we found that SCIN neurogenesis in mouse occurs essentially between E10.5 and E14.5, with a peak around E13.5, and that this time-course is unchanged in *Chat-cKO* mice (Fig. 2b). Furthermore, the birthdate of SCINs predicts their final compartmental localization in the adult mouse striatum, with early-born SCINs (E10.5-E11.5) preferentially populating the striosomes and late-born (E12.5-E14.5) preferentially ending up in the matrix (Fig. 2c), similarly to the rat. Interestingly, *Tshz3* deletion in SCINs profoundly alters this ontogenetic process, with the late-born SCINs now populating the striosomes and the matrix in similar proportions (Fig. 2c). Overall, these findings show that *Tshz3* deletion does not affect the production of SCINs but significantly shifts the settling of late-born SCINs towards the striosomal compartment.

**Fig. 2.**
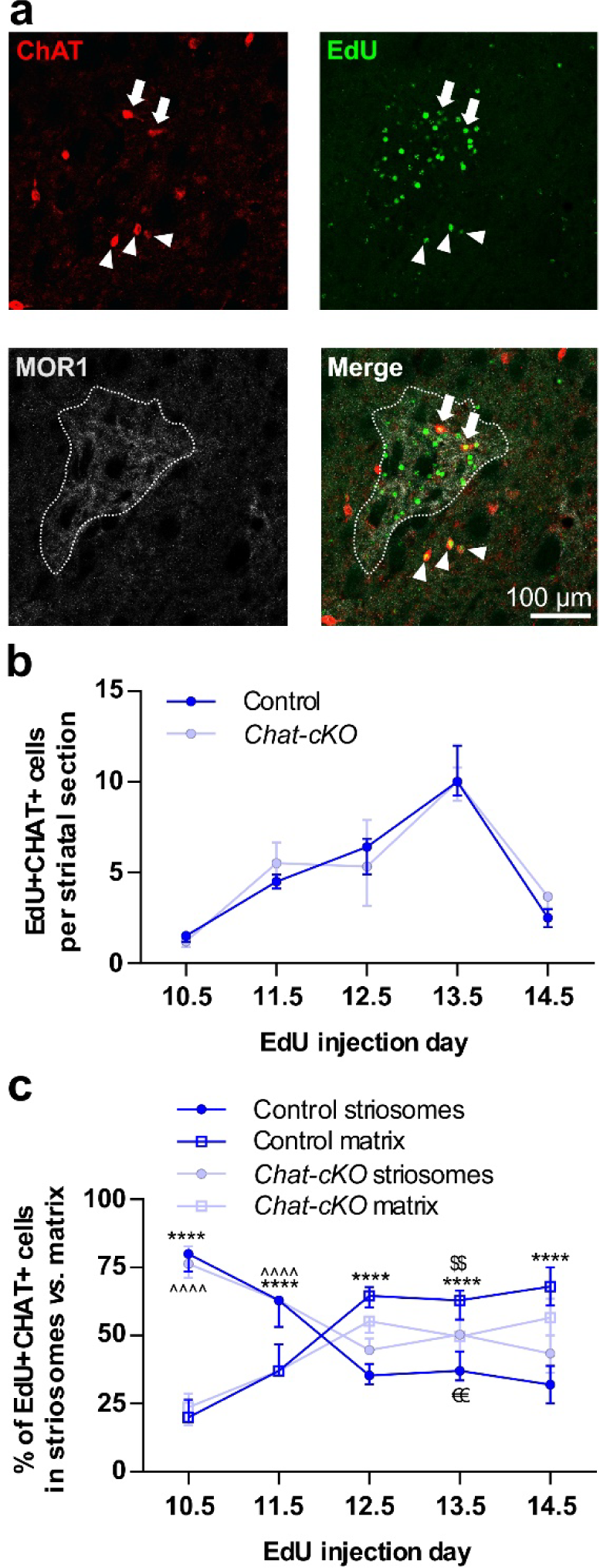
Altered relationship between the birthdates of SCINs and their compartmental distribution in *Chat-cKO* mice. **a**, Illustration of the striosome-matrix distribution of birth-dated SCINs in the adult striatum showed by triple immunodetection of ChAT (red), EdU (green) and MOR1 (light gray). Images are from a *Chat-cKO* mouse after EdU injection in the mother at E13.5. Arrows design EdU-positive/ChAT-positive SCINs residing in the MOR1-positive striosome (dotted line) and arrowheads those located in the surrounding matrix. **b**, Time-course of SCIN neurogenesis expressed as the number of EdU-positive/ChAT-positive cells counted per striatal section after EdU injection at one of the indicated embryonic time points. Note the lack of difference between *Chat-cKO* and control mice, both showing very low SCIN production at E10.5 and E14.5, and a peak at E13.5 (Two-way ANOVA: *F*_genotype_(1,27) = 0.02887, *P* = 0.8663). **c**, Percentages of birth-dated SCINs in the striosomes and matrix. The compartmental distribution of SCINs generated at E10.5 and E11.5 is almost identical in control and *Chat-cKO* mice, with the majority of these cells localized in striosomes. In contrast, while SCINs generated at E12.5-14.5 are found in a significantly larger proportion in the matrix of control mice, their distribution in *Chat-cKO* is around 50% for each compartment. In addition, the percentages of SCINs generated at E13.5 found in striosomes and matrix significantly differ between the two genotypes. Two-way ANOVA: *F*_genotype_(3,54) = 12.39, *P* < 0.0001; *F*_interaction_(12,54) = 44.59, *P* < 0.0001; Tukey’s multiple comparisons post-test: ^^^^*P* < 0.0001, control: striosomes *vs*. matrix; \*\*\*\**P* < 0.0001, *Chat-cKO*: striosomes *vs*. matrix; ^€€^*P* < 0.01, striosomes: control *vs*. *Chat-cKO*; ^$$^*P* < 0.01, matrix: control *vs*. *Chat-cKO*. Data in **b** and **c** are expressed as median ± interquartile range (note that some error bars are shorter than the symbol’s height); they were obtained from 4 mice per genotype for each time point, except at E14.5 (2 control and 3 *Chat-cKO* mice).

### Enrichment of SCINs in the striosomal compartment of *Chat-cKO* mice

In agreement with the above results, counts of SCINs at different antero-posterior levels of the adult striatum do not show significant differences between *Chat-cKO* and control mice (Fig. 3a-c). The total striatal area and the area occupied by MOR1-positive striosomes do not differ either between the two genotypes (Fig. 4a-d). Regarding the distribution of SCINs in control mice, we found that 25.2-33.5% of them localize in the striosome compartment, which represents only 5-10% of the striatal surface, indicating higher SCIN density in striosomes compared to the matrix (Fig. 5a,b). Interestingly, in *Chat-cKO* mice, SCINs are significantly further enriched in striosomes compared to control, reaching ∼40% of the total population (Fig. 5a,b). No difference in the density of striosomal SSPNs, identified by BCL11B immunostaining (Arlotta et al., 2008), was observed between control and *Chat-cKO* mice (Fig. 5c,d). Altogether, these data indicate that *Tshz3* loss specifically affects the compartmental distribution of SCINs by enriching their proportion in the striosomes *vs*. the matrix.

**Fig. 3.**
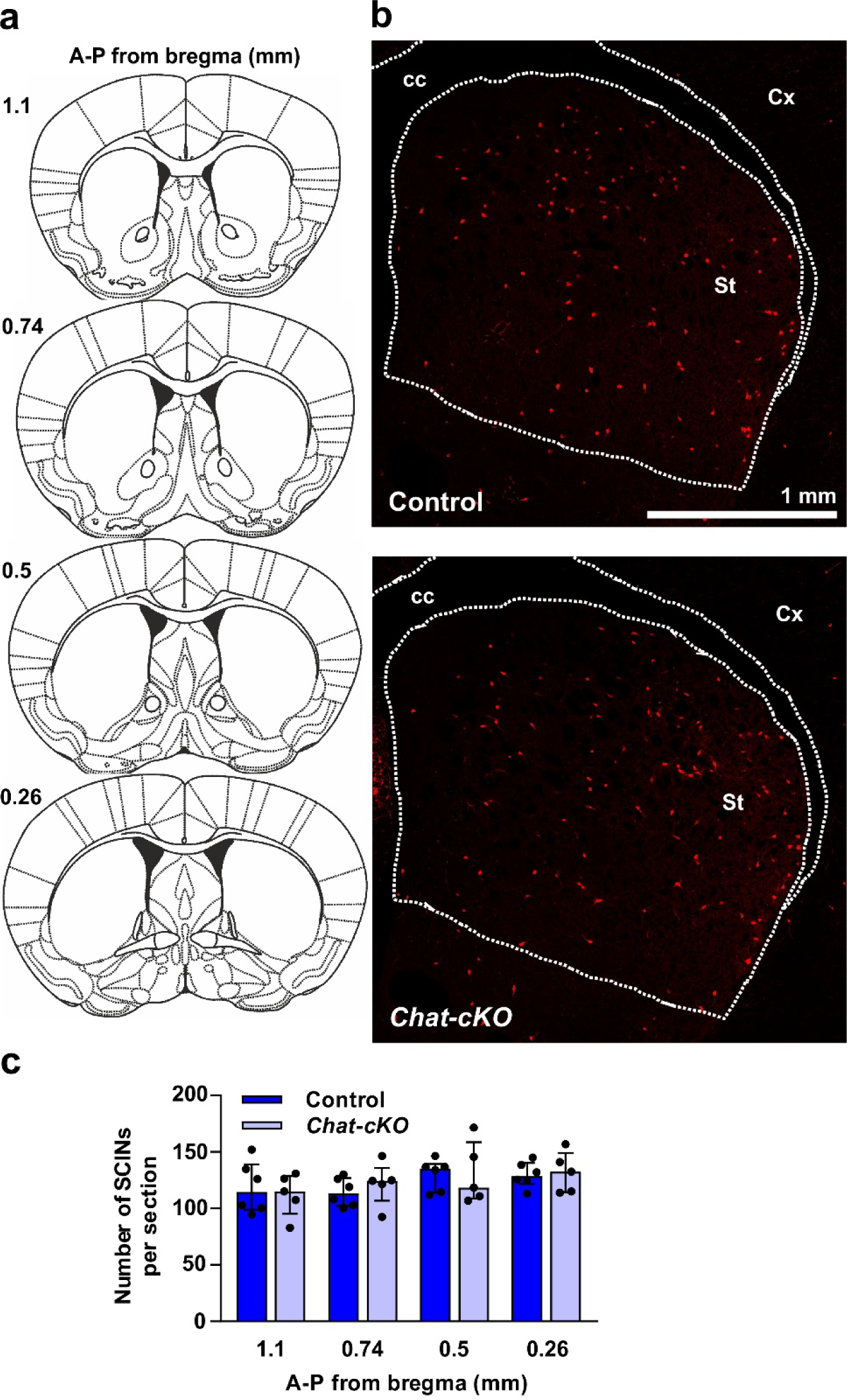
Unchanged numbers of SCINs in *Chat-cKO* mice. **a**, Schemes of coronal mouse brain sections spanning from bregma 1.1 to 0.26 mm, corresponding to the examined antero-posterior (A-P) levels. **b**, Coronal brain sections around 0.5 mm from bregma showing tdTomato-positive (red) SCINs in the striatum of a control and a *Chat-cKO* mouse (abbreviations: cc = *corpus callosum*; Cx = cortex; St = striatum; dotted lines define these structures). **c**, Number of SCINs identified by tdTomato fluorescence per striatal section in control (n = 6 mice) and *Chat-cKO* mice (n = 5 mice). The values do not differ between the two genotypes and are homogeneous at the four A-P levels analyzed. Two-way ANOVA: *F*_genotype_(1,36) = 0.038, *P* = 0.846; *F_A-P_*(3,36) = 2.072, *P* = 0.121; *F*_interaction_(3,36) = 0.249, *P* = 0.862. Data are expressed as median ± interquartile range; dots represent individual values.

**Fig. 4.**
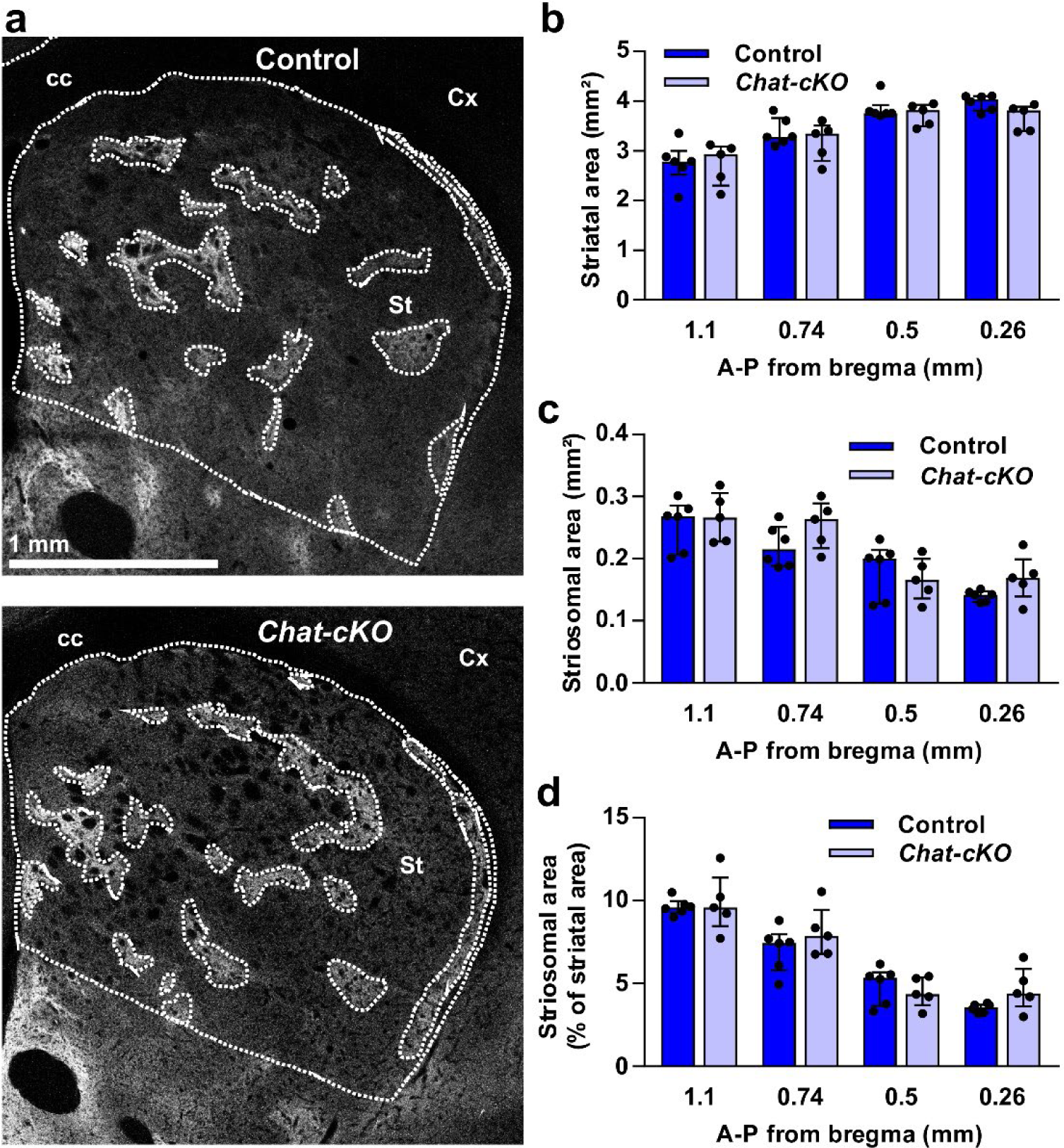
Lack of change in the area of the striatum and of the striosomal compartment in *Chat-cKO* mice. **a**, Coronal brain sections from a control and a *Chat-cKO* mouse showing MOR1-positive (light gray) striosomes surrounded by the MOR1-negative matrix (abbreviations: cc = *corpus callosum*; Cx = cortex; St = striatum; dotted lines define the striatum and MOR1-positive striosomes). **b-d**, Quantitative analyses of the striatal area (**b**), the area of the striosome compartment (**c**) and the striosomal area expressed as percent of striatal area (**d**). No significant difference in any of these parameters is found between control (n = 6) and *Chat-cKO* (n = 5) mice at the four antero-posterior (A-P) levels examined. Two-way ANOVA: *F*_genotype_(1,36) = 2.704, *P* = 0.1088 (**b**); *F*_genotype_(1,36) = 1.936, *P* = 0.1726 (**c**); *F*_genotype_(1,36) = 1.1937, *P* = 0.1726 (**d**). Data are expressed as median ± interquartile range; dots represent individual values.

**Fig. 5.**
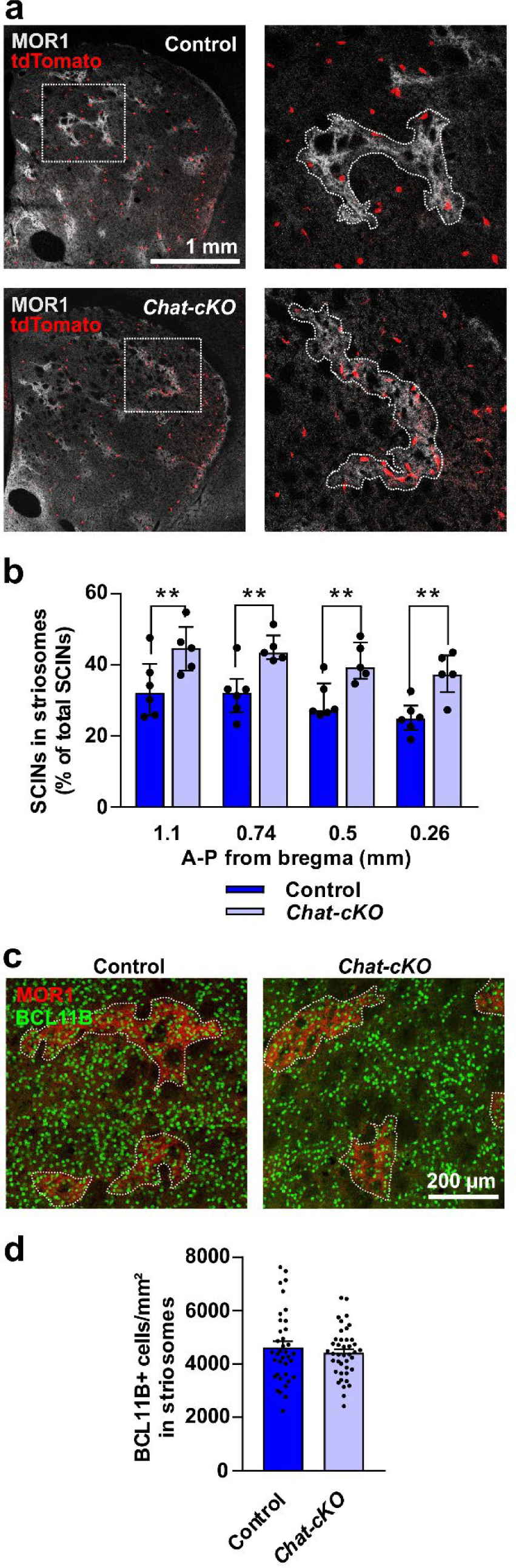
Enrichment of SCINs, but not of SSPNs, in the striosome compartment of *Chat-cKO* mice. **a**, Coronal brain sections from a control and a *Chat-cKO* mouse showing the distribution of tdTomato-positive SCINs (red) in the MOR1-positive striosomes (light gray; dotted lines) and in the MOR1-negative matrix. Higher magnification images of the selected areas (dotted squares) are shown on the right. Note the enrichment of SCINs in the striosomes of *Chat-cKO vs*. control mouse. **b**, Percentage of SCINs per striatal section localized in striosomes. Values are significantly higher in *Chat-cKO* mice (n = 5) compared to control (n = 6) at all antero-posterior (A-P) levels. Two-way ANOVA: *F*_genotype_(1,36) = 38.34, *P* < 0.0001; uncorrected Fisher’s LSD post-test: \*\**P* < 0.01. Data are expressed as median ± interquartile range; dots represent SCIN counts for each mouse. **c**, High-magnification images of the striatum of a control and a *Chat-cKO* mouse showing MOR1-positive striosomes (red) and BCL11B-positive SSPNs (green). **d**, The density of BCL11B-positive SSPNs in the striosome compartment is similar in control and *Chat-cKO* mice (Student’s *t*-test: *P* = 0.424). Counts of BCL11B-positive cells were done in striosomal areas delineated on 19 sections (spanning from bregma 0.26 to 1.10 mm) from 3 mice per genotype. 37 areas from control and 41 *Chat-cKO* mice were analyzed, representing a total counted surface of 0.831 mm^2^ and 0.781 mm^2^, respectively. Data are expressed as mean + SEM; dots represent the SSPN density value of each striosomal area.

### Increased proportion of SCINs with low-frequency, irregular *vs*. high-frequency, regular firing in *Chat-cKO* mice

We next examined whether the above changes are associated to functional modifications in SCINs. To address this issue, we studied their electrophysiological properties in acute brain slices of control and *Chat-cKO* mice, where the tdTomato fluorescence makes them easily recognizable from the other striatal neurons. Compared to controls, SCINs from *Chat-cKO* mice fire action potentials (APs) at significantly lower frequency (Fig. 6a) and less regularly (Fig. 6b; regularity is expressed as inter-spike intervals (ISI) variance), as we reported previously (Caubit et al., 2022). They also produce a significantly smaller voltage sag ratio (VSR) (Fig. 6c) and corresponding hyperpolarization-induced *I*_h_ current (Fig. 6d), which is known to be gated by HCN channels and suppressed by their blocker ZD7288 (Wilson, 2005; Zhao et al., 2016), as verified here (Fig. 6d). Input resistance (42 control SCINs: 161.4 ± 8.3 MΩ; 43 *Chat-cKO* SCINs: 159.4 ± 8.1 MΩ; linear regression: *F*(1,909) = 0.031, *P* = 0.86) and resting membrane potential at the steady state (42 control SCINs: 49.1 ± 1.9 mV; 43 *Chat-cKO* SCINs: 46.3 ± 1.6 mV; Student’s *t*-test: *P* = 0.26,) are similar between the two genotypes (Fig. 6e). SCIN spontaneous firing pattern and frequency are variable and have been classically used to define them as regular-, irregular- and burst-firing both on slice and *in vivo* (Bennett and Wilson, 1999; Wilson et al., 1990). Accordingly, we found a wide range of both ISI variance (regularity) and firing frequency, although we never observed any SCIN of the burst-firing subtype. Interestingly, we measured a strong inverse correlation between AP firing frequency and ISI variance in both control and *Chat-cKO* SCINs (Spearman r > 0.8; Fig. 6f), while the other pair combinations of the three parameters examined (AP frequency, ISI interval and VSR) have a weak correlation (r < 0.4; not shown). We thus established a cutoff corresponding to the median ISI variance value measured in control SCINs (i.e., 1179) in order to univocally classify these neurons as either high-frequency, regular firing (hfrSCINs), or low-frequency, irregular firing (lfiSCINs), so as hfrSCINs show an ISI variance smaller and lfiSCINs greater than the cutoff value (Fig. 6f). Representative examples of the electrophysiological properties of a hfrSCIN and an lfiSCIN from a control mouse are illustrated in Fig. 6g. We next examined whether these two subpopulations are differentially affected by the conditional *Tshz3* loss by calculating the ratio between the number of lfiSCINs over the total number of SCINs recorded from each mouse (Fig. 6h). This ratio shows that in control mice lfiSCINs are the minority (0.39), while in *Chat-cKO* mice they are the majority (0.58), which accounts for the overall decrease of spontaneous firing frequency and regularity in *Chat-cKO* mice.

**Fig. 6.**
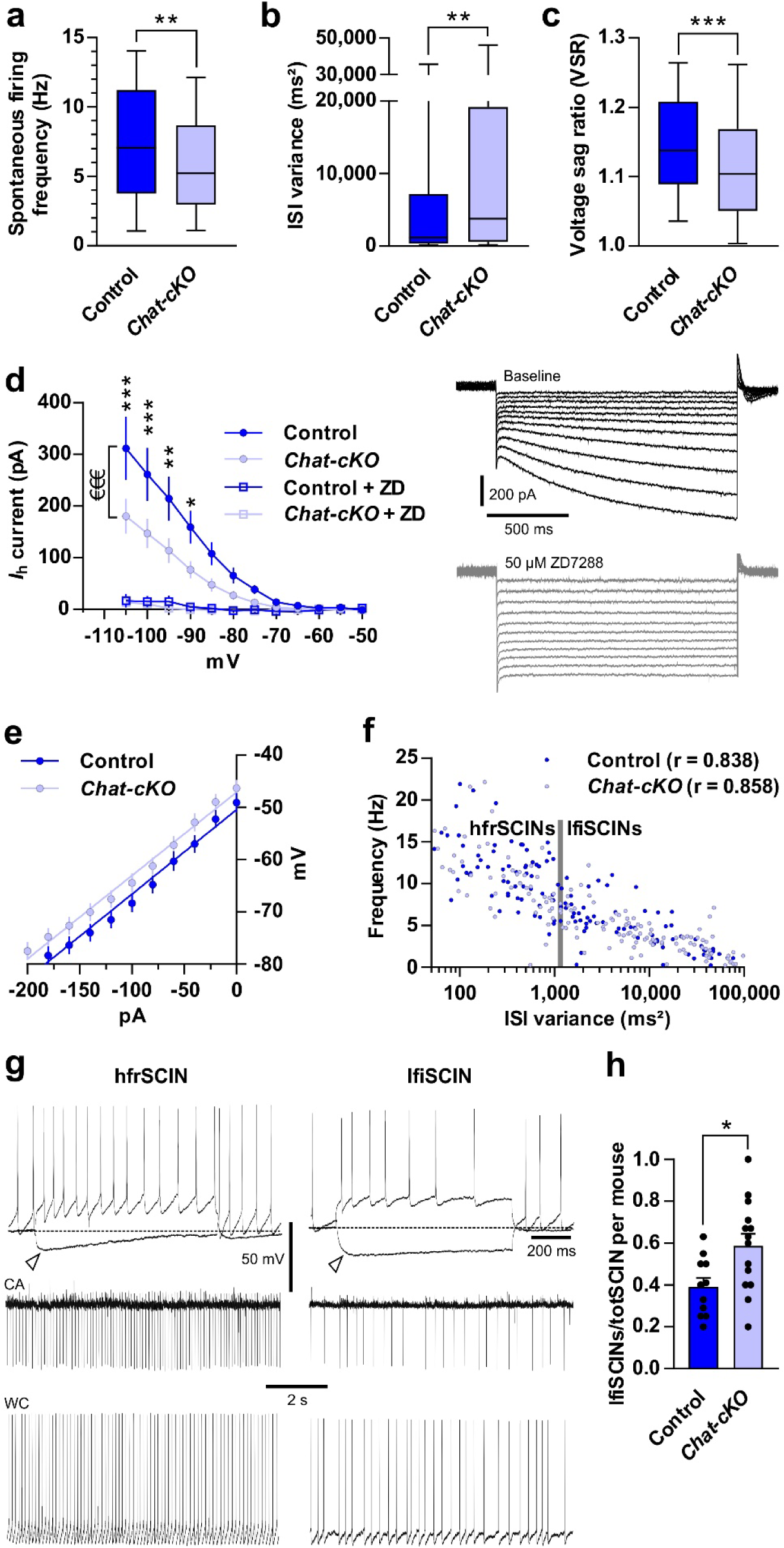
Changes in the firing features of SCINs in *Chat-cKO* mice. **a**, The frequency of spontaneous action potential firing is significantly reduced in *Chat-cKO* SCINs compared to control (control: 129 SCINs; *Chat-cKO*: 149 SCINs; \*\**P* = 0.008, Mann-Whitney test). **b**, The ISI variance is increased *Chat-cKO* SCINs, indicating a less regular activity (control: 128 SCINs; *Chat-cKO*: 143 SCINs; \*\**P* = 0.008, Mann-Whitney test). **c**, The voltage sag ratio (VSR) is significantly decreased in *Chat-cKO* SCINs (control: 163 SCINs; *Chat-cKO*: 206 SCINs; \*\*\**P* = 0.0004, Mann-Whitney test). **d**, The *I*_h_ current is reduced in *Chat-cKO* (control: 12 SCINs; *Chat-cKO*: 11 SCINs; ^€€€^Two-way ANOVA: *F*(1,252) = 23.5; uncorrected Fisher’s post-test: \**P* < 0.05, \*\**P* < 0.01, \*\*\**P* < 0.001). Note the drastic reduction of the *I*_h_ by bath application of the HCN channel blocker ZD7288 (ZD). Traces show an example of the currents evoked by hyperpolarizing voltage pulses (from - 55 to −105 mV by −5 mV steps, holding potential −50 mV) before (black) and after (gray) ZD7288 application in a control SCIN: note the suppression of the early component (sag) due to the *I*_h_ and the reduction of the late component. **e**, The graph shows the current-voltage relationships obtained from 41 control and 42 *Chat-cKO* SCINs, and the respective linear best fits to calculate input resistance, which is similar between the two genotypes (see Results). **f**, Plot of the ISI variance-frequency showing their correlation and the Spearman r values of control and *Chat-cKO* SCINs (control: 127 SCINs; *Chat-cKO*: 150 SCINs; *P* < 0.0001 for both). The vertical gray bar represents the control median value (1,179 ms²) used as cutoff for classifying SCINs as hfrSCIN (≤ 1,179) and lfiSCIN (> 1,179). **g**, Sample traces showing the representative electrophysiological features of a hfrSCIN (left) and an lfiSCIN (right) from a control mouse. Upper row: responses to a depolarizing and a hyperpolarizing current step (+180 and –120 pA, respectively); arrowheads show the voltage sag (hfrSCIN, VSR = 1.6; lfiSCIN, VSR = 1.29). Lower rows: spontaneous firing in cell-attached (CA) and whole-cell (WC) configuration: note the higher frequency and regularity of action potential discharge of the hfrSCIN (11.7 Hz, ISI variance = 284.7 ms²) compared to the lfiSCIN (3.1 Hz, ISI variance = 12,189 ms²). **h**, The proportion of lfiSCINs over the total number of SCINs (totSCINs) recorded from each mouse is significantly higher in *Chat-cKO* (n = 94 SCINs from 14 mice) compared to control mice (n = 83 SCINs from 11 mice) (\**P* < 0.05, Mann-Whitney test). Data in **a-c** are expressed as 25th-75th percentiles (boxes), 5th-95th percentiles (whiskers) and median (horizontal line); in **d**, **e**, **h** as mean ± SEM; in **f** as single SCIN values; dots in **h** represent the ratio value of each mouse.

### Enrichment of SCINs with low-frequency, irregular firing in the striosomes of *Chat-cKO* mice

Having shown that lfiSCINs are the predominant subtype in *Chat-cKO* mice, we next studied how hfrSCINs and lfiSCINs are distributed across the striosome-matrix compartments and whether this is affected by *Tshz3* loss. For doing so, some of the recorded SCINs were injected with biocytin via the patch-clamp pipette in order to localize them on MOR1-immunostained sections (Fig. 7a). This approach shows that hfrSCINs and lfiSCINs have similar compartmental distributions in control animals (Fig. 7b,c), with about 20% of both populations lying in the striosomes. In *Chat-cKO* mice, the striosome-matrix distribution of hfrSCINs is not significantly modified (Fig. 7b), whereas the proportion of lfiSCINs in the striosomes is dramatically increased *vs*. control (52.4% *vs*. 21.1%; Fig. 7c). Altogether, these findings clearly show that the loss of *Tshz3* produces a drastic shift of SCIN electrophysiological features towards the lfiSCIN subtype, which selectively enriches the striosome compartment. This results in an overall slower firing rate of striosomal *vs*. matrix SCINs in *Chat-cKO* mice (Fig. 7d).

**Fig. 7.**
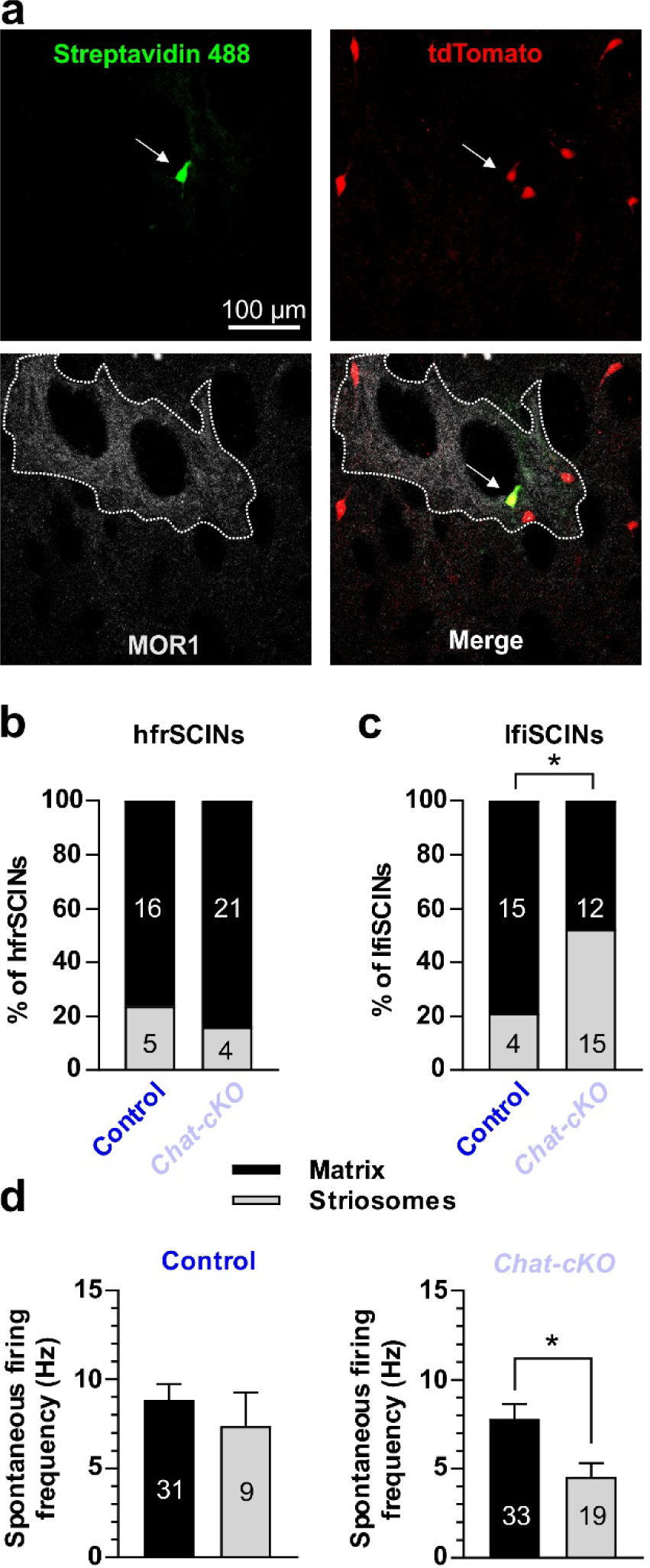
Enrichment of SCINs with low frequency/irregular firing in the striosomes of *Chat-cKO* mice. **a**, Images showing a recorded SCIN (arrow) injected with biocytin and revealed by both streptavidin 488 (green) and tdTomato (red) fluorescence (upper row). This neuron is localized within a MOR1-positive (light gray) striosome (lower row), outlined by the dotted line. **b**,**c**, Distribution of hfrSCINs (**b**) and of lfiSCINs (**c**) within the striosome and the matrix compartments. hfrSCINs are similarly distributed in the two genotypes (χ² = 0.4423, *P* = 0.506), while lfiSCINs are enriched in the striosomes of *Chat-cKO vs*. control mice (χ² = 5.476, \**P* = 0.019). **d**, Average spontaneous firing frequency of all identified SCINs in either striosomes or matrix. There is no compartmental difference in control mice, while in *Chat-cKO* mice striosomal SCINs fire at a significantly lower frequency. Data in **b-d** are obtained from 5 control and 7 *Chat-cKO* mice; the numbers of recorded and identified SCINs are reported in the columns. Data in **d** are expressed as mean + SEM; \**P* = 0.011, Student’s *t*-test.

### Reduced levels of striatal VAChT predominating in the striosomes of *Chat-cKO* mice

To further characterize the consequences of *Tshz3* deletion, we measured the expression levels of VAChT, the protein responsible for acetylcholine loading into synaptic vesicles, which is considered as a key regulator of cholinergic transmission (Prado et al., 2013). Since VAChT level does not vary in heterozygous *Chat-Cre* mice compared to control mice (Chen et al., 2018), changes in *Chat-cKO* mice would reflect modified cholinergic activity due to *Tshz3* deletion. Consistent with the reduced firing of SCINs shown in Fig. 6a,h, we observed a significant decrease of VAChT levels determined by western blot analysis (Fig. 8a,b) and of VAChT immunofluorescent staining in the striatum of *Chat-cKO* mice (Fig. 8c). We then examined the compartmental distribution of VAChT-positive fibers using EphA4 receptor immunostaining, which marks the matrix (Fig. 8d).

**Fig. 8.**
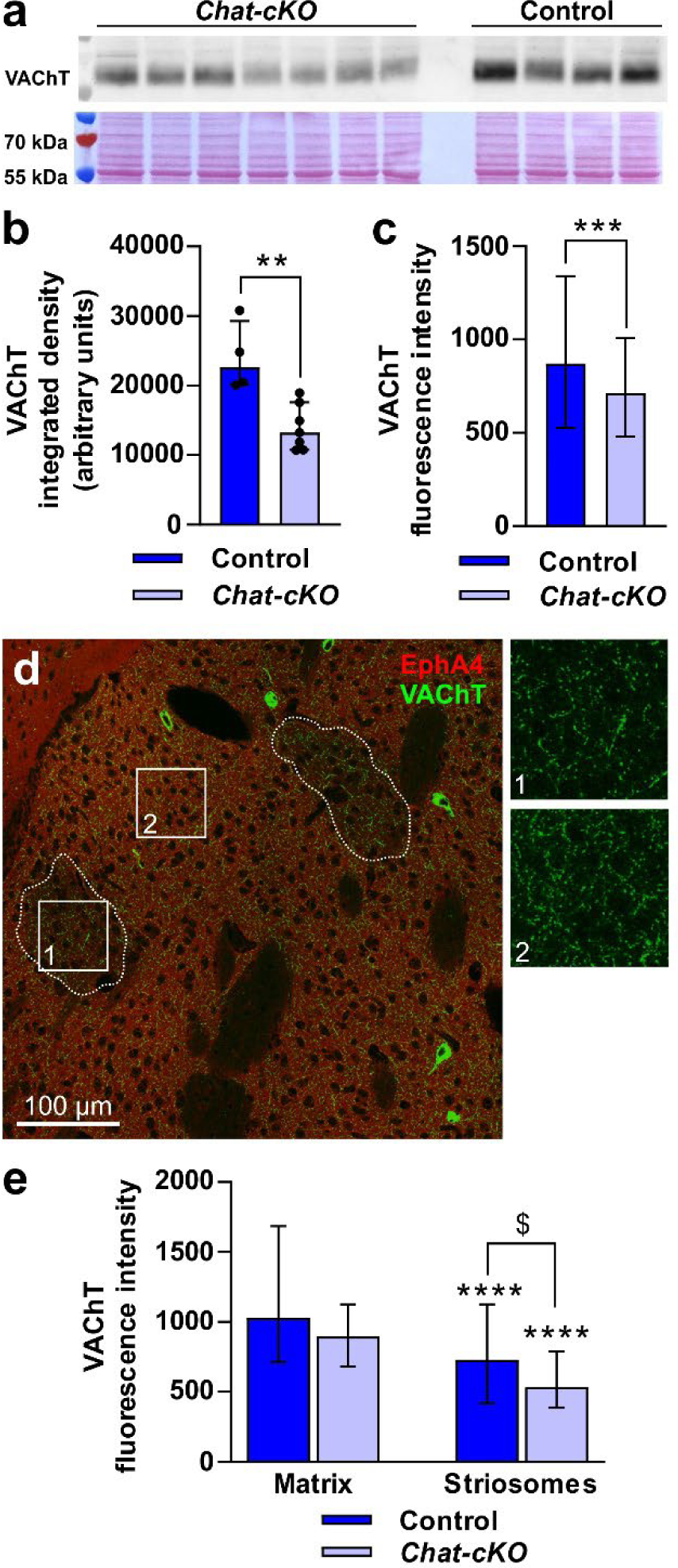
Reduced VAChT expression in *Chat-cKO* mice. **a**, **b**, Western blot analysis of striatal VAChT. **a**, Immunoblot of the VAChT protein (upper panel) and corresponding total protein detection by Ponceau red staining (lower panel). **b**, Integrated density histogram of VAChT showing a significant decrease in *Chat-cKO* mice compared to control (\*\**P* < 0.01, Mann-Whitney test). **c**, The intensity of VAChT immunofluorescence is significantly lower in the striatum of *Chat-cKO* mice compared to control (\*\*\**P* < 0.001, Mann-Whitney test). **d**, Coronal brain section of a control mouse striatum showing the EphA4-positive matrix (red) that allows identifying two EphA4-negative striosomes (dotted lines), merged with VAChT immunostaining (green) labeling the cholinergic neuropil. Note the lower VAChT signal in the striosome compartment compared to the matrix (square 1 and 2, respectively; squares are zoomed on the right). **e**, VAChT expression is significantly lower in the striosomes *vs*. the matrix in both control and *Chat-cKO* mice, and significantly reduced in the striosomes of *Chat-cKO* mice compared to controls (Kruskal-Wallis ANOVA, *P* < 0.0001; Dunn’s post-test: ^$^*P* < 0.05; \*\*\*\**P* < 0.0001 *vs*. matrix, same genotype). Data are expressed as median ± interquartile range and were obtained from 6 control and 6 *Chat-cKO* mice. A total of 110 ROIs per compartment were measured in control and 114 in *Chat-cKO* mice.

In control animals, the intensity of VAChT labeling is lower in the striosomes than in the matrix (Fig. 8e). This shows that the cholinergic neuropil of the striatum is compartmentalized in the mouse, as previously reported in the human (Hirsch et al., 1989) as well as in the macaque and the cat (Graybiel et al., 1986), based on the observation that ChAT-positive fibers are sparse in the striosomes while dense in the matrix delineated by acetylcholinesterase staining. In *Chat-cKO* mice, the levels of VAChT in the striosomes are also significantly lower than in the matrix, and they are reduced compared to control striosomal values (Fig. 8e). A trend towards reduction is also observed in the matrix of *Chat-cKO vs*. control mice, but the difference did not reach significance. Collectively, these data indicate that cholinergic activity is lower in the striosomes compared to the matrix in both control and *Chat-cKO* mice. Furthermore, consistent with the depressed firing activity of SCINs, they suggest that *Tshz3* loss in SCINs results in reduced striatal cholinergic tone and that this reduction is more pronounced in the striosomal compartment.

## DISCUSSION

Here we describe the temporal profile of SCIN neurogenesis in the mouse in relation to their striosome-matrix distribution in the adult striatum, and demonstrate a prolonged commitment of late-born SCINs to populate the striosomes in *Chat-cKO* mice. *Tshz3* deletion in SCINs also results in decreased frequency and regularity of their spontaneous AP firing, and more specifically in an enrichment of lfiSCINs over hfrSCINs, which in addition concentrate into striosomes. Moreover, levels of VAChT are decreased, especially in the striosomal compartment. Overall, these defects in SCIN striosome-matrix distribution and electrophysiological properties may profoundly affect striatal microcircuitry by altering cholinergic tone and may play a role in the stereotyped behaviors that characterize our *Chat-cKO* mouse model of ASD.

SCINs are an heterogeneous neuronal population due to their multiple embryonic origins (MGE, septal neuroepithelium and preoptic area) and the expression of different gene sets during early brain development (Ahmed et al., 2019; Knowles et al., 2021; Lopes et al., 2012; Marin et al., 2000). Studies in the rat have shown that the birthdates of both SCINs (van Vulpen and van der Kooy, 1998) and SSPNs (Graybiel and Hickey, 1982; Song and Harlan, 1994; van der Kooy and Fishell, 1987) predict their compartmental localization in the adult striatum, with striosomal neurons generated earlier than those of the matrix. In the mouse, this process is conserved for SSPNs, with early- and late-born neurons preferentially populating the striosomes and the matrix, respectively (Lebouc et al., 2020). Concerning SCINs, it was known that early-born (before E12.5) populate the lateral region of the adult striatum, while late-born (after E12.5) the medial (Chen et al., 2010), but the exact contribution of these two migration waves to the striosome-matrix compartments remained unknown. This is an important issue considering the different circuitry and function of these two compartments (Brimblecombe and Cragg, 2017; Crittenden and Graybiel, 2011; Crittenden et al., 2017), and the role of SCINs in the striatum as key modulators of synaptic transmission/plasticity and neuronal excitability (Abudukeyoumu et al., 2019; Poppi et al., 2021; Prado et al., 2017). Here, we confirm that SCINs are born mainly between E11.5 and E14.5 (Chen et al., 2010) with a production peak at E13.5. We further show that their final compartmental localization follows a pattern similar to that of SSPNs, namely with early-born SCINs populating the striosomes and late-born the matrix, and that the transition between the two phases of this program takes place between E11.5 and E12.5. Another question is the relative proportion of the overall population of SCINs laying in either compartment at adulthood. We found that ∼30% of SCINs are localized in MOR1-positive striosomes, although a previous paper, using different methodology to define striosomes and identify SCINs, mentioned that these interneurons populate the striosomes only “occasionally” (Crittenden et al., 2014). Given our data showing that striosomes occupy 5-10% of the total striatal surface, which agrees with previous reports from rat and mouse (Johnston et al., 1990; Kubota and Kawaguchi, 1993; Lee et al., 2008), this indicates that the density of SCINs in the two compartments is uneven, being much higher in the striosomes. This result is comparable to previous data showing that the density of SCINs in striosomes, identified as enkephalin-rich patches, is twice that of the matrix in the dorsal caudate nucleus of the cat (Martone et al., 1994), but differs from a study in the rat reporting that less than 12% of SCINs are located in the striosomal compartment, defined as calbindin D_28k_-poor patches (Kubota and Kawaguchi, 1993). Whether these discrepancies are due to species differences and/or to the markers used to define the striosome-matrix compartments or the striatal region analysed remains to be determined.

SCINs are also heterogeneous from an electrophysiological point of view: they show autonomous AP firing with a wide range of frequencies and patterns, and exhibit the nonspecific *I*_h_ cation current mediated by HCN channels whose amplitude is also variable (Bennett et al., 2000; Bennett and Wilson, 1999; Caubit et al., 2022; Goldberg and Wilson, 2010; Kawaguchi, 1993; Lozovaya et al., 2018). Our study provides a quantitative analysis of SCINs’ firing patterns, allowing to classify them as “high-frequency, regular firing” (hfrSCINs) and “low-frequency, irregular firing” (lfiSCINs) and to define their respective proportions. This analysis also shows that SCINs’ firing rate is inversely correlated with its regularity, confirming literature data (Bennett and Wilson, 1999) and further extending them on a larger sample. This is an interesting point, since it can allow classifying SCINs on the basis of either parameter, opening new perspectives for the study of SCIN electrophysiological properties and how they are affected by different experimental conditions. Thank to this classification, we show that the striosomes and the matrix contain similar proportions of hfrSCINs and lfiSCINs, indicating that the two compartments do not differ in the functional properties of their populating SCINs. Therefore, taking into account the ontogenesis of SCINs, it can be suggested that the birthdate of these interneurons does not predict their electrophysiological features.

Cholinergic transmission abnormalities have been reported in the brain of ASD patients (Marotta et al., 2020) and in the striatum of mice modeling ASD (Karvat and Kimchi, 2014; Rapanelli et al., 2017b), as well as in the striatum of rodents with drug-induced stereotyped behaviors (Aliane et al., 2011; Athnaiel et al., 2022; Crittenden et al., 2017). Increasing evidence also point to abnormal striatal compartmentation and striosome-matrix imbalance as a pathological signature of ASD and/or stereotyped behaviors (Crittenden and Graybiel, 2011; Ferhat et al., 2023; Kuo and Liu, 2017, 2020; Murray et al., 2015). Furthermore, literature data link psychostimulant-induced stereotyped behaviors to, on one hand, striosome activation measured as *Fos*-*Jun* gene family expression (Canales and Graybiel, 2000; Saka et al., 2004; Sako et al., 2010), and, on the other hand, to striatal cholinergic signaling (Aliane et al., 2011; Crittenden et al., 2014; Crittenden et al., 2017; Peter et al., 2017). However, whether and how SCIN dysfunction relates to abnormal compartmentation has not been investigated in the context of ASD. Here we address this compelling issue using our *Chat-cKO* mouse model of ASD-related stereotyped behaviors. We provide evidence that, in this model, the *Chat-Cre* becomes active around E14.5, so that the deletion of *Tshz3* in immature SCINs should occur from this time point onwards. Therefore, changes in SCINs should not happen during their neurogenesis, which peaks at E13.5, but rather during their late differentiation process, their migration from the proliferative zones to the future striatal area and/or their final positioning and connectivity within the striatal network (Fragkouli et al., 2009; Knowles et al., 2021; Magno et al., 2017). Accordingly, we show that the time-course of SCIN neurogenesis is almost identical in control and *Chat-cKO* mice. In addition, the total number of SCINs in the adult striatum is unaffected in *Chat-cKO* mice, suggesting that *Tshz3* loss after E14.5 does not affect either the viability of these interneurons, which is postnatally regulated by local inputs from SSPNs (Sreenivasan et al., 2022). However, interestingly, *Tshz3* loss perturbs the striosome-matrix compartmental distribution of SCINs in adult mice, with an increased percentage of them localized in the striosomes, reaching more than 40% of the total population. As the areas of the striosomal compartment and of the whole striatum are unaffected, the striosomal SCIN density is thus drastically enhanced. This increase can be attributed to changes in the developmental trajectory of the late-born SCINs, which, instead of preferring the matrix as in control, end up in the striosomes and the matrix with similar proportions. In contrast, the density of striosomal SSPN is not affected, underlining the SCIN-specificity of these distributional changes. Overall, these data show that *Tshz3* deletion in SCINs leads to subtle rearrangements of the striosome-matrix distribution of these interneurons, rather than to major rearrangements of the striosomal area as observed in other ASD models (Ferhat et al., 2023; Kuo and Liu, 2017).

Our electrophysiological analysis of SCINs from *Chat-cKO* mice shows a marked decrease of their spontaneous firing frequency and regularity, as well as reduced *I*_h_ current amplitude. This is consistent with the literature reporting that both these parameters of spontaneous firing are proportional to the average amplitude of the *I*_h_ current and the expression of HCN channels (Bennett et al., 2000; Cheng et al., 2019; Choi et al., 2020; Zhao et al., 2016). Interestingly, *Cntnap2* KO mice, in which this gene is deleted in SCINs and in other neuronal populations and which exhibit abnormal behaviors mimicking the core ASD features, show similar modifications in SCIN firing (Ahmed et al., 2023). In *Chat-cKO* mice, the decrease of SCIN discharge activity could result in a reduced acetylcholine tone in the striatum and, consistently, we measured a decrease in VAChT expression. As the total number of SCINs is unchanged, such a decrease is unlikely to be due to a reduction of the cholinergic neuropil, but rather to a reduced release of acetylcholine linked to the lower neuronal activity. In line with these findings, we previously reported a significantly reduced expression of the *Slc18a3* gene coding for VAChT in another mouse model with *Tshz3* deletion in SCINs that also shows stereotypies (see Supplementary Table 1 in (Caubit et al., 2021)). This further supports the view that the stereotyped behavior associated with *Tshz3* deletion might be linked to decreased acetylcholine release. Such decreased cholinergic tone could lead to changes of SSPN excitability, which is mainly mediated by M1 muscarinic receptors, and could perturb striatal synaptic transmission and plasticity, which are mediated by muscarinic and nicotinic receptors at both pre- and postsynaptic level (Abudukeyoumu et al., 2019; Bonsi et al., 2011; Calabresi et al., 2000; Gertler et al., 2008).

The overall decrease in SCINs firing rate and regularity observed here translates into a higher proportion of lfiSCINs over the total number of recorded SCINs. Interestingly, the higher striosomal density of SCINs in these mice is due to a specific enrichment of lfiSCINs in this compartment, leading to an overall decrease of SCIN firing in the striosomes compared to the matrix. Therefore, *Chat-cKO* mice show not only altered compartmental distribution of SCINs but also different functional properties of the SCINs populating the striosomes and the matrix, as also supported by the decreased VAChT expression in striosomes. This could lead to an imbalanced functioning of the two compartments, perturbing the mutual regulation of excitability and synaptic transmission between SCINs and the other striatal neurons (Abudukeyoumu et al., 2019; Ahmed et al., 2019; Crittenden et al., 2017; Inoue et al., 2016).

Overall, the findings of the present study provide evidence that ASD-related stereotyped behaviors are not necessarily linked to gross developmental and anatomo-functional rearrangements of the striosome-matrix compartments, but can be generated by very subtle changes in the compartmental distribution and function of a subpopulation of SCINs. Whether or not such SCIN abnormalities are a hallmark of stereotypies in other models of ASD and/or other pathological conditions characterized by stereotyped behaviors is an interesting issue that would deserve further investigation. While designating SCINs as a target for alleviating stereotyped behaviors, our work raises several questions and technical challenges, since it suggests that the best strategy should be to specifically target striosomal SCINs, and in particular lfiSCINs. This would require a thorough molecular analysis of SCIN subpopulations allowing to distinguish and manipulate them, ultimately leading to the identification of druggable molecular targets.

## Contributions

J.M., J.G., P.S. and X.C. performed the histological experiments and the quantitative analyses. J.M. and P.G. performed the patch-clamp experiments and analyzed the electrophysiological data. J.M., J.G. and X.C. generated and maintained the transgenic mouse line. F.C. performed western blot experiments. J.M., J.G., X.C. and A.F. performed mouse genotyping. L.K.-L.G., L.F., X.C. and P.G. conceived the project, supervised the work and wrote the paper. All authors have read and approved the final manuscript.

## Acknowledgments

This work received support from the French government under the Programme “Investissements d’Avenir”, Initiative d’Excellence d’Aix-Marseille Université via A*Midex funding (NeuroMarseille Institute, AMX-19-IET-004; MarMaRa Institute, AMX-19-IET-007), the French National Research Agency (ANR) “TSHZ3inASD” project grant n°ANR-17-CE16-0030-01 (to L.F. and L.K.-L.G.), the Fédération pour la Recherche sur le Cerveau (to L.F.), the Centre National de la Recherche Scientifique (CNRS), and Aix-Marseille University. J.M. and J.G. were supported by PhD grants from the Ministère de l’Enseignement Supérieur, de la Recherche et de l’Innovation. We wish to thank the IBDM imaging platform (France-BioImaging/PiCsL infrastructure (ANR-10-INSB-04-01)) and mouse facility.

## METHODS

### Mouse strains and genotyping

The *Tshz3^+/lacZ^*, *Tshz3^flox/flox^*, *Chat-Cre* and *Ai14* (*Rosa26-STOP-tdTomato*) mouse lines have been described previously (Caubit et al., 2016; Caubit et al., 2008; Chabbert et al., 2019; Madisen et al., 2010; Mao et al., 1999; Rossi et al., 2011). *Chat-Cre* mice crossed with *Ai14* (*Chat-Cre;Ai14*) mice were used as control, and henceforth referred to as control. Male heterozygous *Chat-Cre;Tshz3^flox/flox^* mice were crossed with female *Tshz3^flox/flox^*;*Ai14* to generate *Chat-cKO;Ai14* mice, and hereafter referred to as *Chat-cKO.* Thus, cholinergic cells (including SCINs) of both control and *Chat-cKO* mice express the tdTomato red fluorescent marker. Animals were genotyped as described previously (Caubit et al., 2022; Chabbert et al., 2019). Experimental procedures were in agreement with the recommendations of the European Communities Council Directive (2010/63/EU). They have been approved by the “Comité d’Ethique en Expérimentation Animale de Marseille C2EA-14” and the project authorization delivered by the French Ministry of Higher Education, Research and Innovation (ID numbers 57-07112012, 2019020811238253-V2 #19022, 2020031615241974-V5 #25232 and 2022042118061902-V4 #36912). No randomization was used and no animals or samples were excluded from the different analyses performed.

### Electrophysiology

Electrophysiological data were obtained from 30 *Chat-cKO* and 27 control male mice, aged between 21 and 28 days. Procedures were similar to those described previously (Caubit et al., 2022). Briefly, acute coronal slices (250 µm-thick) containing the cortex and the striatum were cut using a S1000 Vibratome (Leica) in ice-cold solution containing (in mM): 110 choline, 2.5 KCl, 1.25 NaH_2_PO_4_, 7 MgCl_2_, 0.5 CaCl_2_, 25 NaHCO_3_, 7 glucose, pH 7.4. Slices were kept at room temperature in oxygenated artificial cerebrospinal fluid (ACSF), whose composition was (in mM): 126 NaCl, 2.5 KCl, 1.2 MgCl_2_, 1.2 NaH_2_PO_4_, 2.4 CaCl_2_, 11 glucose and 25 NaHCO_3_, pH 7.4. Electrophysiological recordings were performed in ACSF at 34-35 °C and flowing at ∼2 ml/min. SCINs of the dorsal striatum were identified by infrared and fluorescence video microscopy (due to their tdTomato fluorescence), and by their electrophysiological properties (Bennett et al., 2000; Bennett and Wilson, 1999; Kawaguchi et al., 1995). They were recorded first in cell-attached and then in whole-cell configuration using patch-clamp borosilicate micropipettes (5-6 MΩ) filled with an internal solution containing (in mM): 125 K-gluconate, 10 NaCl, 1 CaCl_2_, 2 MgCl_2_, 0.5 BAPTA, 19 HEPES, 0.3 Na-GTP, and 1 Mg-ATP, pH 7.3. Electrophysiological data were acquired by an AxoPatch 200B amplifier and pClamp 10.7 software (Molecular Devices, Wokingham, UK). The current-voltage (I-V) relationship was obtained in voltage-clamp mode from a subset of SCINs, by measuring the membrane response at the end of current steps from −200 to −20 pA (20 pA steps lasting 800 ms), and input resistance was calculated by linear regression analysis, i.e. as the slope of the linear best fit of the I-V relationship of each recorded neuron. The voltage sag ratio (VSR) was calculated from the response to a −120 pA current step as the peak voltage drop (sag) against the voltage at the end of the current pulse (Haghdoust et al., 2007; Maisano et al., 2012). Such relatively small current step was chosen because, with larger steps, the sag amplitude was extremely variable between different SCINs. *I*_h_ current was measured at peak from the response to voltage from −55 to −105 mV (−5 mV steps lasting 800 ms) at a holding potential of −50 mV. Spontaneous action potential (AP) firing was analyzed in terms of discharge frequency, expressed in Hz, and regularity, expressed as the variance of inter-spike intervals (ISI). Note that spontaneous AP firing was analyzed only from cell-attached recordings, which were done before switching to whole-cell. In some cases, spontaneous firing was not detectable in cell-attached configuration, and in other cases whole-cell parameters (VSR, I-V relationship) were discarded due to unstable recording. For these reasons, the number of samples may vary between different data sets. In some cases, SCINs were injected with biocytin (5 mg/ml) via the patch-clamp pipette to allow localizing them in relationship with the striosome-matrix compartments (see below for staining procedure).

### Fluorescence immunostaining and histology

#### Primary polyclonal antibodies

rabbit anti-µ_1_ opioid receptor (MOR1, 1:1000, Abcam, ab10275); goat anti-choline acetyltransferase (ChAT, 1:100, Millipore, ab144P); rabbit anti-vesicular acetylcholine transporter (VAChT, 1:1000, SYSY, 139103); guinea pig anti-vesicular acetylcholine transporter (VAChT, 1:500, SYSY, 139105); chick anti-ß-Galactosidase (ß-Gal, 1:1000, Abcam, ab9361); rat anti-B-cell lymphoma/leukemia 11B (BCL11B/CTIP2) (1:1000, Abcam, ab18465); goat anti-ephrin-A4 (EphA4, 1:150, R&D systems, AF641); guinea-pig anti-TSHZ3 (1:2000; (Caubit et al., 2008)).

#### Secondary antibodies

donkey anti-chick Alexa Fluor 488, donkey anti-guinea pig Alexa Fluor 488, donkey anti-rabbit Alexa Fluor 647, donkey anti-rabbit Alexa Fluor 546, donkey anti-rat Alexa Fluor 488 (Invitrogen), and donkey anti-goat Cy3 (Jackson ImmunoResearch Laboratories).

For studies on the embryonic expression of *Tshz3* and cholinergic markers, wild-type, *Chat-Cre*;*Ai14* and *Tshz3^+/lacZ^* embryos were externalized, their brains were quickly dissected and fixed by immersion for 2 h in 4% PFA, except those of *Tshz3^+/lacZ^* embryos that were fixed overnight in 4% PFA. All brains were cryoprotected overnight, embedded in OCT and stored at −80 °C until cryosectioning at 16 or 20 μm. For immunostaining, the cryosections were blocked in 3% BSA and PBST (phosphate buffer saline (PBS) with 0.1% Triton X-100) for 1 h at room temperature (RT) and then incubated overnight at 4 °C in primary antibodies diluted in the same solution. After washing with PBS (three times; 10 min), the sections were incubated in secondary antibodies 2 h at RT. For some experiments, sections were counterstained for 5 min in 0.1 µg/ml DAPI solution (ThermoFisher).

Postnatal immunostaining studies were performed on mice aged 28-32 days. Animals were anesthetized by buprenorphine (0.1 mg/kg, i.p.) and ketamine + xylazine (100 + 10 mg/kg, respectively, i.p.), and trans-cardially perfused with 4% paraformaldehyde (PFA) in 0.1 M phosphate buffer (PB). Brains were removed and post-fixed in 4% PFA for at least 24 h, cryoprotected in PBS with 30% sucrose for 24 h and kept at −80 °C until use for cryostat sectioning (40 µm-thick; Leica, CM 3050S). Sections were kept in cryoprotectant solution (20% glycerol, 30% ethylene glycol in 0.02 M PB at pH 7.4) at −20 °C until immunostaining. For immunostaining, free floating sections were washed with PBS and blocked with 1% bovine serum albumin (BSA) in PBST for 1 h at RT. They were then incubated in primary antibodies diluted in blocking solution (PBST with 1% BSA) overnight at 4 °C, then washed with PBS three times and incubated 2 h at RT in secondary antibodies diluted 1:1000 in blocking solution.

The numbers of animals used for the different experiments are the following: for striosomal and striatal surface, SCIN counting and striosome-matrix compartmental localization of SCINs and SSPNs: n = 6 control and 5 *Chat-cKO* mice; for VAChT expression: n = 6 control and 6 *Chat-cKO* mice. For biocytin staining, slices were fixed in 4% PFA (overnight) immediately after electrophysiological recordings, then washed three times in PBS and kept in cryoprotectant solution at −20 °C. For localizing the recorded SCINs in the striosome or matrix compartment, slices were incubated 2 h at RT with streptavidin Alexa Fluor 488 (1:500) and rabbit anti-MOR1 antibodies, then processed with donkey anti-rabbit Alexa Fluor 647 secondary antibodies.

In all cases, the sections were washed and mounted on Superfrost Plus slides, and coverslipped with Immu-Mount^TM^ (Thermo Scientific) for imaging on a laser scanning confocal microscope (Zeiss LSM780 with Quasar detection module). Spectral detection bandwidths (nm) were set at 415-473 for DAPI, 498-561 for GFP, 570-630 for Cy3 and 657-758 for 647; pinhole was set to 1 Airy unit.

### EdU birth-dating and immunolabeling

For EdU (5-ethynyl-2′-deoxyuridine) birth-dating assays, pregnant mice were injected intraperitoneally with EdU at 25 µg/g of body weight at either embryonic day (E) 10.5, E11.5, E12.5, E13.5 or E14.5. For each time point, 2 mice were injected. Afterwards, 18 control and 19 *Chat-cKO* adult offspring mice were trans-cardially perfused and their brains cut in 40 µm-thick cryostat sections as above. EdU staining was performed using Click-iT EdU Imaging Kit (C10337, Invitrogen) following the manufacturer’s protocol. In brief, sections were first incubated in Click-iT EdU reaction components for 30 minutes to stain EdU by Alexa Fluor 488 azide, then washed twice with PBST. Sections were then incubated in primary goat anti-ChAT and rabbit anti-MOR1 antibodies diluted in blocking solution (PBST with 1% BSA) overnight at 4 °C, then washed three times in PBS and incubated for 2 h at 4 °C in donkey anti-rabbit Alexa Fluor 647 and donkey anti-goat Cy3 secondary antibodies diluted in blocking solution. Finally, sections were washed, mounted and imaged as described below.

### Image analysis

Unbiased counting of neurons and surface measurements were done on confocal images of coronal brain sections using ImageJ (National Institutes of Health open-source software). They were performed blind to the genotype in the dorsal striatum, bilaterally.

For birth-dating experiments, counts of EdU-positive/ChAT-positive cells were performed on 6 sections/mouse and averaged. For SCIN counting and striosome-matrix surface characterization, measurements were performed at four antero-posterior striatal levels, corresponding to 1.1, 0.74, 0.5 and 0.26 mm from bregma (Paxinos and Franklin, 2001). For each level, 2 sections were analyzed and the values averaged per animal. For each level, the whole area of the striatum was scanned and reconstructed by joining the acquired fields in the x–y axis using a tiling option in the ZEN acquisition software (Zeiss). A line from the bottom of the lateral ventricle to the piriform cortex was drawn to separate the dorsal striatum from the ventral striatum (mainly the nucleus accumbens), which was excluded from the analysis (e.g., Fig. 3b and 4a). MOR1-positive striosomes were selected with the “wand” tool and their area was measured with the “measure” option. Areas were obtained as pixels and then converted in mm².

Counts of tdTomato-labeled cells (i.e., SCINs) were performed both in the whole dorsal striatum and in the striosome compartment using ImageJ “image calculator” and “analyze particles” tools. BCL11B-positive cells (i.e., SSPNs) were counted in striosomes only. Cells at the border of the two compartments were considered as striosomal.

Quantification of VAChT fluorescence signal in striatal compartments was performed on 0.72 mm^2^ images (resolution: 11.37 pixels/mm) that were acquired in 16-bit range using a 40X/1.4 oil objective and with zoom factor = 0.7. Acquisitions were performed for VAChT in channel S1 (488 nm, laser intensity: 1.2%) and for EphA4 in channel S2 (561 nm, laser intensity 5%) at 3.14 μs pixel dwell time. For VAChT signal, laser intensity and photomultiplier tube gain was set so as to occupy the full dynamic range of the detector. On each image, two regions of interest (ROIs) of 4857 µm^2^ were positioned in the striosomal compartment, delineated by the lack of EphA4 staining, and two in the neighboring matrix, and the mean fluorescence intensity per ROI was measured semi-automatically using Fiji ImageJ from NIH (https://imagej.net/Fiji/Downloads). An adjusted background value per image, determined as the mean of the signal measured in 10 myelin bundles, was subtracted from the ROI values. A total of 110 ROIs per compartment were measured in control and 114 in *Chat-cKO* mice, with a minimum of 6 images analyzed from 3-4 sections per mouse.

### Western blotting

Experiments were performed on 7 *Chat-cKO* and 4 control male mice aged 25-30 days. Animals were sacrificed by decapitation, the brains quickly removed and immediately frozen at −80 °C. In a cryostat set at −20 °C, tissue samples of the dorsal striatum were collected from each brain hemisphere using a trocar/cannula system (1 mm diameter), weighted and stored frozen at −80 °C until use. For immunoblotting, samples were homogenized at 4 °C with a Potter homogenizer using 50 µl of modified radioimmunoprecipitation assay (RIPA) buffer per 1 mg of tissue. The RIPA buffer (50 mM Tris-HCl at pH 7.4, 300 mM NaCl, 1 mM EDTA, 1% NP40, and 0.5% sodium deoxycholate) was supplemented with a protease inhibitor cocktail (complete EDTA free, Roche, Basel, Switzerland). Homogenates were centrifuged at 12000 g for 10 min at 4 °C. Proteins present in supernatant were solubilized with Laemmli sample buffer and 40 µl of each sample were resolved by NuPAGE 4-12% Bis-Tris gel (MES buffer, Thermo Scientific, WG1401) and transferred onto nitrocellulose membranes. Membranes were stained with Ponceau S to visualize protein loading and VAChT was revealed with rabbit anti-VAChT (1:1000, SYSY, 139103) by ECL procedure (Clarity Western ECL, Biorad) on a chemiluminescence imager (ChemiDoc, Biorad). For illustration purposes, image editing was performed using ImageJ software (http://rsb.info.nih.gov.insb.bib.cnrs.fr/ij/) and was limited to linear brightness/contrast adjustment. Quantification of the immunoblot bands was performed using ImageJ. Results are the integrative intensities minus background.

### Statistical analysis

Statistical analysis was performed by Prism 7.05 (GraphPad Software, USA). Before statistical analysis, data sets were tested for normality by the D’Agostino & Pearson’s normality test. When normality was not achieved by at least one data set, they were analyzed using non-parametric tests, otherwise they were analyzed by parametric tests. Two-tailed Student’s *t*-test or Mann-Whitney test were used for comparing two data sets. Kruskal-Wallis (ANOVA) test followed by Dunn’s multiple comparisons test was used to compare 3 or more data sets. Two-way ANOVA followed by a multiple comparisons post-test (uncorrected Fisher’s least significant difference post-test, or Tukey’s post-test) was used to analyze the influence of 2 categorical variables. Spearman’s r was used to calculate correlation: the degree of correlation was considered “weak” when r < 0.4, “moderate” when 0.4 ≤ r ≤ 0.8 and “very strong” when r > 0.8 (Akoglu, 2018). Data are expressed as mean ± SEM when they have a normal distribution; when their distribution does not pass the normality test, they are expressed as box & whiskers (25th-75th & 5th-95th percentiles, respectively; horizontal bar = median), or as median ± interquartile range. Each sample value is represented by a black dot, except when the number of samples is too high to be clearly represented so. The significance threshold is set at *P* < 0.05. The tests used, *P* values and sample sizes [as number (n) of recorded neurons, mice or histological sections] are indicated in the Methods, Figures and/or Results.

